# Regulatory network controlling tumor-promoting inflammation in human cancers

**DOI:** 10.1101/352062

**Authors:** Zhe Ji, Lizhi He, Aviv Regev, Kevin Struhl

## Abstract

Using an inducible, inflammatory model of breast cellular transformation, we describe the transcriptional regulatory network mediated by STAT3, NF-κB, and AP-1 factors on a genomic scale. These regulators form transcriptional complexes that directly regulate the expression of hundreds of genes in oncogenic pathways via a positive feedback loop. This inflammatory feedback loop, which functions to various extents in many types of cancer cells and patient tumors, is the basis for an “inflammation” index that defines cancer types by functional criteria. We identify a network of non-inflammatory genes whose expression is well correlated with the cancer inflammatory index. Conversely, the inflammation index is negatively correlated with expression of genes involved in DNA metabolism, and transformation is associated with genome instability. Inflammatory tumors are preferentially associated with infiltrating immune cells that might be recruited to the site of the tumor via inflammatory molecules produced by the cancer cells.

## INTRODUCTION

Tumor-promoting inflammation is a hallmark of cancer and plays important regulatory roles during cell transformation, invasion, metastasis and treatment resistance (Coussens and Werb, 2002; Mantovani et al., 2008; Grivennikov et al., 2010). An inflammatory reaction during tumor development occurs in near all solid malignancies (de Martel and Franceschi, 2009). Pre-existing inflammation promotes subsequent cancer development, accounting for 15%~20% of cancer deaths (Mantovani et al., 2008). Tumor-promoting inflammation also plays critical roles in immunosuppression (Grivennikov et al., 2010). For example, STAT3 promotes the expression of PD-L1 and PD-L2 in cancer cells, suppressing immune cell activity (Chen and Han, 2015).

Tumor-associated inflammation results from both an intrinsic pathway, in which mutations in cancer cells activate inflammatory gene expression, and an extrinsic pathway, in which cytokines and chemokines secreted by tumor-associated immune cells create inflammatory microenvironments (Mantovani et al., 2008). Oncogenes can trigger a gene expression cascade, resulting in activation or overexpression of pro-inflammatory transcription factors such as NF-κB, STAT3 and AP-1, with the resulting production of cytokines and chemokines (Coussens and Werb, 2002; Iliopoulos et al., 2009; Grivennikov et al., 2010). For example, NF-κB signaling pathway is activated upon activation of oncogenes RAS and MYC, through activation of IL-1β (Shchors et al., 2006; Guerra et al., 2007; Iliopoulos et al., 2009), and the proto-oncoprotein Src kinase can directly phosphorylate and activate STAT3 (Turkson et al., 1998).

In previous work, we have described an inducible model of cellular transformation in which transient activation of v-Src oncoprotein converts a non-transformed breast epithelial cell line into a stably transformed state within 24 hours (Iliopoulos et al., 2009; Hirsch et al., 2010). The stably transformed cells form foci and colonies in soft agar, show increased motility and invasion, form mammospheres, and confer tumor formation in mouse xenografts (Iliopoulos et al., 2009; Hirsch et al., 2010). This epigenetic switch between stable non-transformed and transformed states is mediated by an inflammatory positive feedback loop involving NF-κB and STAT3 (Iliopoulos et al., 2009; Iliopoulos et al., 2010). By integrating motif analysis of DNase hypersensitive regions with transcriptional profiling, we found that > 40 transcription factors are important for transformation and identified putative target sites directly bound by these factors (Ji et al., 2018).

A few transcriptional regulatory circuits involved in this transformation model have been identified, and these are important in some other cancer cell types and human cancers (Iliopoulos et al., 2009; Iliopoulos et al., 2010; Iliopoulos et al., 2011b; Polytarchou et al., 2012). During transformation, STAT3 acts through pre-existing nucleosome-depleted regions bound by FOS, and expression of several AP-1 factors is altered in a STAT3-dependent manner (Fleming et al., 2015). However, the connection between STAT3, NF-κB, and AP-1 factors as well as the underlying transcriptional regulatory circuits has not been described on the whole-genome level.

Here, we define the transcriptional network mediated by the combined action of NF-κB, STAT3 and AP-1 factors (JUN, JUNB and FOS) on a genomic scale in this breast transformation model. In contrast to previous studies (Fleming et al., 2015; Ji et al., 2018), this network is defined by genes that are induced during transformation by the binding of NF-κB, STAT3 and AP-1 factors to common target sites either through their cognate motifs or via protein-protein interactions. Based on this common NF-κB, STAT3 and AP-1 network, we develop a “cancer inflammation index” to define cancer types, both in cell lines and in patients, by functional criteria. As this inflammation index is based on the common regulatory network, it is distinct from and more specific than indices based simply on gene expression profiles that arise from multiple regulatory inputs. In addition, we identify many non-inflammatory genes whose expression is positively or negatively correlated with the cancer inflammation index, leading to the observation the inflammation is linked functionally to other aspects of cancer as well as genomic instability. Lastly, we show that inflammatory tumor samples preferentially contain contaminating immune and stromal cells from the tumor microenvironment, consistent with the idea that immune cells might be recruited to the site of the tumor via inflammatory molecules produced by the cancer cells.

## RESULTS

### Co-localization of STAT3, NF-κB and AP-1 factors on target sites *in vivo*

During transformation, NF-κB, STAT3, and AP-1 (JUN, JUNB and FOS) levels increase in the nucleus (Figure S1A). In accord with previous results using chemical inhibitors and siRNA knockdowns (Iliopoulos et al., 2009), CRISPR-cas9-mediated knockouts of these factors (Figure S1B) cause decreased transformation as assayed by growth under conditions of low attachment (Rotem et al., 2015)(Figure S1C). Our previous ChIP-seq analysis mapped protein-binding sites to nucleosome-depleted regions before and after (24 hours tamoxifen treatment) transformation (Fleming et al., 2015), but did not provide more detailed localization.

The ChIP-seq data reveals many target sites bound by STAT3 (89,764 sites), NF-κB (56,539 sites) and AP-1 factors (95,958 sites for JUNB, 152,443 sites for FOS and 118,245 sites for JUN) (Figures 1A, S1D and S2A). Less than 17% of factor binding sites are located in promoter regions, and the majority of sites are located in other regulatory regions, including ~30% that are in active enhancers (Figure S2A). As expected from the heteromeric nature of AP-1 factors, binding levels of JUN, JUNB and FOS are well correlated with each other (Pearson correlation coefficiencies >= 0.83), which are nearly comparable to biological replicates (Figure S2B). Based on this, we next grouped the AP-1 factors together to learn their binding features on the chromatin. Also as expected, STAT3 binding sites strongly overlap with FOS binding sites (Fleming et al., 2015) and with the other AP-1 factors (89% of STAT3 binding sites) (Figure S3A). In addition, 89% of NF-κB binding sites overlap with AP-1 factors (Figure S3A). 38% of STAT3, NF-κB, and AP-1 sites are located in the same czs-regulatory regions (CRRs), which is far more frequent than expected by chance (*P* < 10^−200^; Chi-squared Test) (Figure S3A). Despite the strong co-localization, STAT3, NF-κB and AP-1 factors can independently bind to specific sites (Figure S3A), and the pairwise correlation values are lower than the correlation values between biological replicates (Figure S3B).

**Figure 1.**
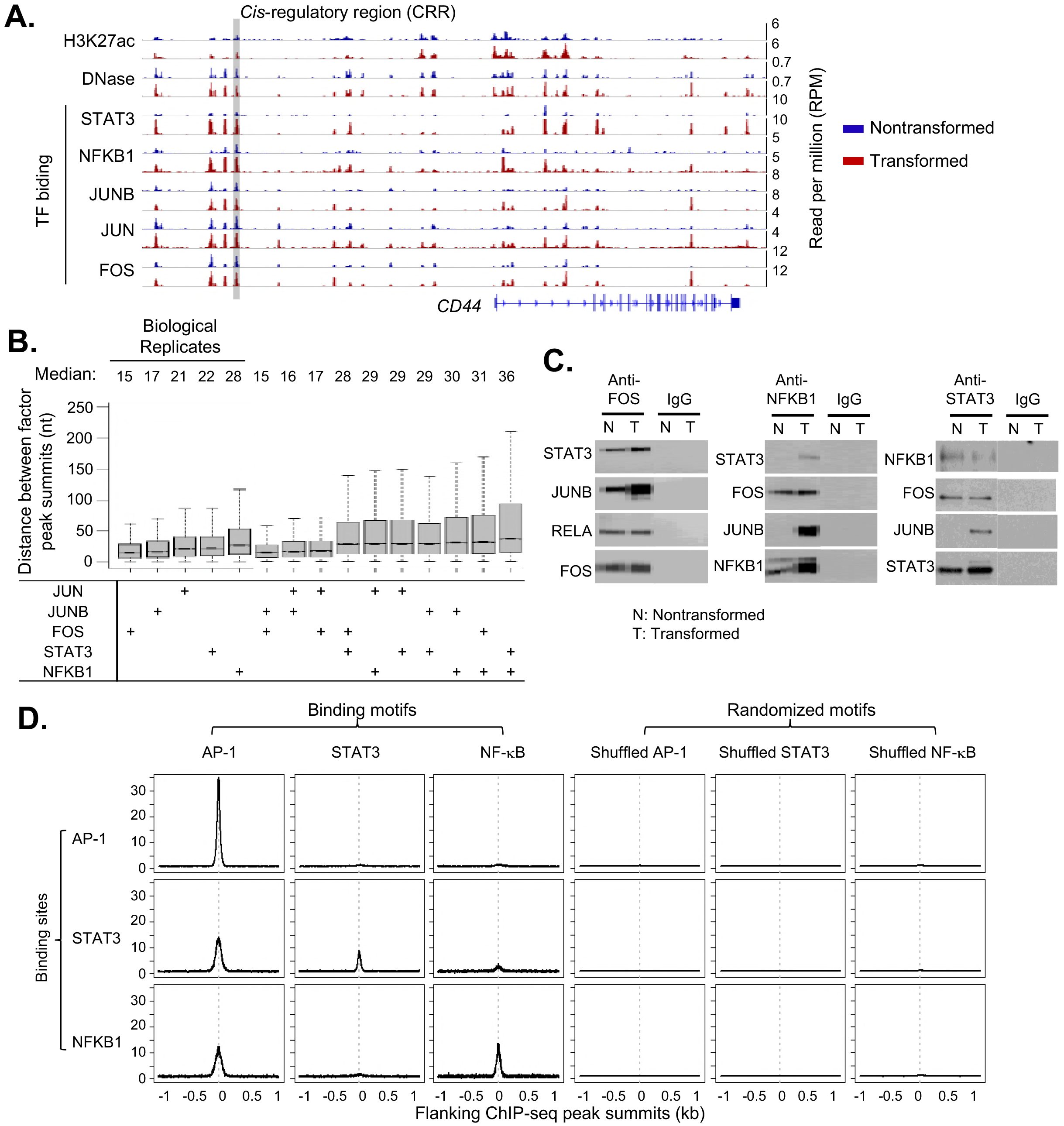
Features of transcription factor binding. (A) An example genomic locus with CD44 gene showing ChIP-seq of STAT3, NFKB1, JUN, JUNB, FOS, and H3K27ac, and DNase-seq to map open chromatin regions before (blue) and after (red) transformation. (B) Distance between peak summits of indicated factors locating in the same *cis*-regulatory region. (C) Co-immunoprecipitation experiment showing the interactions between FOS, JUNB, STAT3 and NF-κB in the nucleus of non-transformed (N) and transformed (T) cells. IgG IP was used as the control. (D) Distribution of AP-1, STAT and NF-κB consensus motif around factor peak summits. As the control, we randomly shuffled AP-1, STAT and NF-κB consensus motifs and plotted the motif distribution around peak summits.

Additional results suggest that STAT3, NF-κB and AP-1 factors co-bind to target sites as a multiprotein complex. First, the median distance of peak summits for all pairwise combinations of AP-1 factors, STAT3, and NF-κB ranges between 15-30 bp, and these values are comparable to those obtained for biological replicates of the relevant individual factors (Figure 1B). Second, co-immunoprecipitation experiments show that STAT3, NF-κB and AP-1 factors interact with each other in the nucleus (Figure 1C).

### Sequence motifs associated with binding of STAT3, NF-κb and AP-1 factors

To address which factors are primarily responsible for binding site specificity, we performed motif analysis on 50 bp sequences around peak summits. As expected, 60% of AP-1 factor binding peaks contain an AP-1 motif (Figure S3C). Interestingly, 38% of STAT3 binding sites and 30% of NFKB1 sites also have an AP-1 motif, while 24% of STAT3 sites have a STAT motif and 16% NFKB1 sites have a NF-κB motif (Figure S3C). Furthermore, the AP-1 motif is well located in STAT3 and NFKB1 peak summits (Figure 1D). We did not find significant co-localization of AP-1, STAT and NF-κB motifs in the same CRRs. These results indicate that a significant fraction of STAT3 and NF-κB binding is mediated through the interaction with AP-1 factors. In contrast, only a small minority of AP-1 binding sites contain a STAT or NF-κB motif (Figures 1D and S3C), and STAT and NF-κB motifs show modest enrichment around AP-1 peak summits (Figure 1D). Thus, binding of AP-1 factors occurs predominantly via interactions with AP-1 motifs, presumably reflect a direct protein-DNA interaction. In contrast, in addition to directly binding via their motifs, STAT3 and NF-κB can also bind to AP-1 motifs, presumably via protein-protein interactions with AP-1 factors. However, we did not observed significant motif differences in AP-1 sites bound by AP-1 factors alone or those co-bound with STAT3 and/or NF-κB.

### STAT3, NF-κb and AP-1 factors co-regulate key genes in various oncogenic pathways

Co-binding of STAT3, NF-κB and AP-1 factors regulates gene expression and chromatin status. Genes up-regulated during transformation (Figure 2A) tend to have increased binding of STAT3, NFKB1, JUN, JUNB and FOS at promoters/enhancers (Figure 2B, D). Furthermore, isogenic cell lines lacking individual factors can block the up-regulation of a subset of genes during transformation (Figure 2C), with the STAT3 knockout showing the most drastic effects. As there are multiple members of the NF-κB and AP-1 families, it is likely that the weaker effect on transcription are due to redundancy among family members. With respect to chromatin structure, regions bound by increasing numbers of these factors tend to have higher accessibility (Figure S4A) and acetylation levels (Figure S4B). In addition, differential binding levels of STAT3, NFKB1 and AP-1 factors during transformation are positively correlated with dynamic chromatin accessibility (Pearson Correlation Coefficient >= 0.44) (Figure S4C), and open chromatin regions bound by more factors tend to show increased accessibility (Figure S4D).

**Figure 2.**
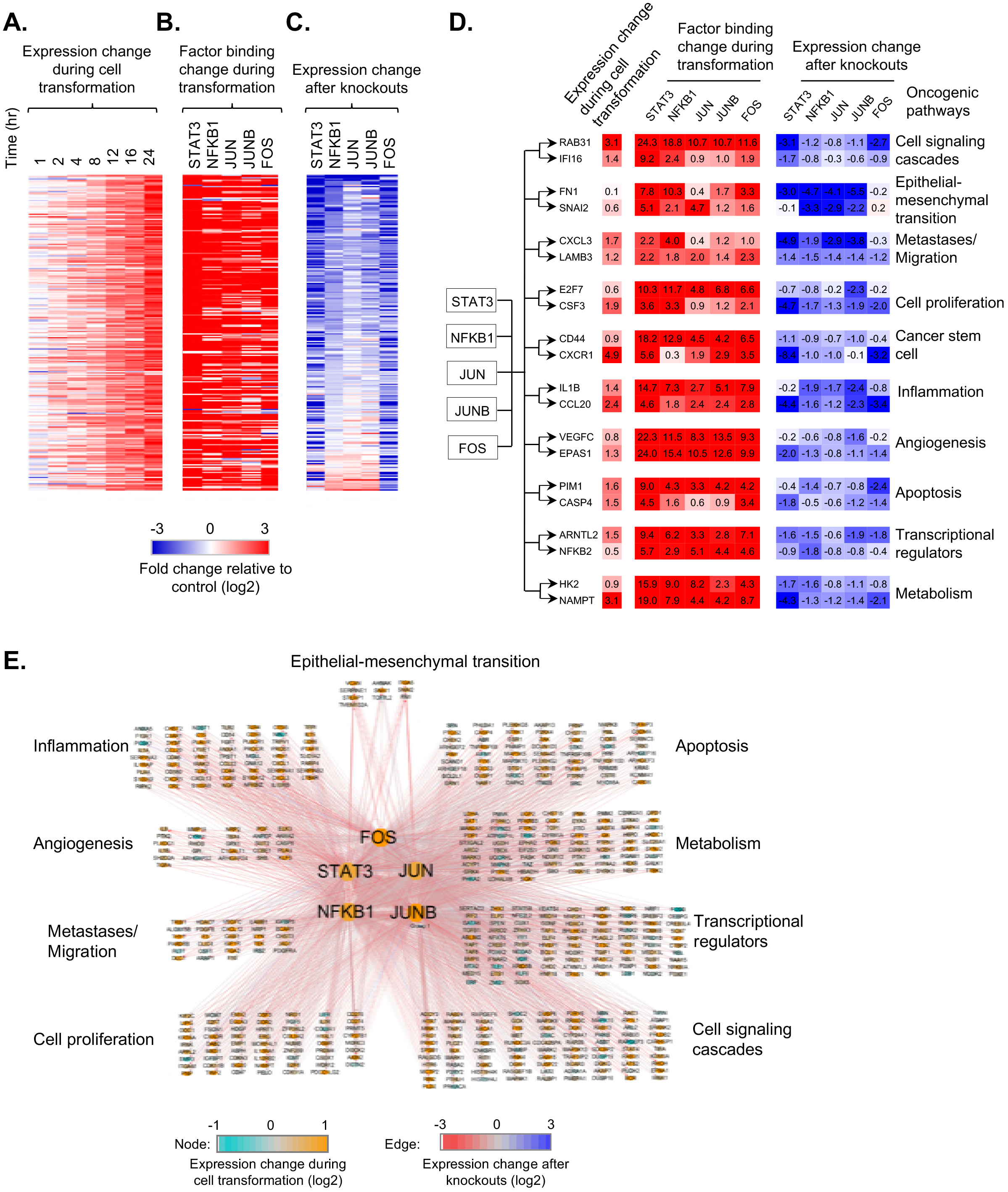
Co-binding and common targets of STAT3, NF-κb and AP-1 factors. (A-C) For each gene showing continuously up-regulation during cell transformation (A), fold-changes of factor binding levels in promoters/enhancers (B) and gene expression changes after factor knockouts at 24hr after the cell transformation (C) are shown. (D) Examples of common target genes of STAT3, NFKB1 and AP-1 factors regulating different oncogenic pathways. The fold-changes of expression during transformation, factor binding level changes in promoters/enhancers and gene expression changes after factor knockout at 24 hr after transformation are shown. The degree of up-regulation (red) and down-regulation (blue) is indicated by the color intensity. (E) Transcriptional network mediated by STAT3, NFKB1 and AP-1 factors in different oncogenic pathways as indicated. The nodes represent factors and target genes in the network. The edges represent direct binding of factors in promoter/enhancer regions of target genes, and gene expression change upon factor knockouts.

We identified 1,461 genes that are common targets of STAT3, NFKB1, JUN, JUNB and FOS and that show increased binding of at least four factors (>1.5 fold) during transformation and downregulation upon at least four factor knockouts (Table S1). These genes are enriched (Benjamini–Hochberg FDR < 0.005) in cancer-related processes (Figure S5), and they include genes involved in cell signaling cascades (e.g. RAB13, IF116, ZAK), inflammatory response (e.g. IL1B, IL1R1, SERPINA1), cell proliferation (e.g. CSF3, E2F7, E2F8) regulation of apoptosis (e.g. PIM1, CARD6, BCL2L1), angiogenesis (e.g. RHOB, CEGFC, EPAS1), and cell migration/metastases (e.g. LAMB3, CXCL3 and PLAU)(Figure 2D, E). Cancer stem cells are generated during the transformation process in our model (Iliopoulos et al., 2011a), and STAT3, NF-κB and AP-1 factors activate key genes (e.g. CD44, CXCR1 and ITGA7) mediating cancer stem cell formation. Cancer metabolism is a key component of oncogenesis (Pavlova and Thompson, 2016), and important metabolic enzyme genes (e.g. HK2 and NAMPT), are also regulated by these factors (Figure 2D, E). Thus, STAT3, NF-κB, and AP-1 are the major factors involved in tumor-promoting inflammation, and they bind to and promote the expression of key genes in various oncogenic pathways (Figure 2E).

### The positive feedback inflammatory loop that promotes transformation is extensive

In previous work, we described a few specific regulatory circuits that constitute an inflammatory positive feedback loop required for transformation in this experimental model (Iliopoulos et al., 2009; Iliopoulos et al., 2010; Fleming et al., 2015). Here, we show that this positive feedback loop is far more extensive. In particular, during transformation, STAT3, NF-κB (NFKB1 and NFKB2) and AP-1 factors (JUN, JUNB, FOS, FOSL1, FOSL2, and ATF3) are transcriptionally up-regulated as are many upstream regulators in the IL6/STAT3, IL1/NF-κB and TNF/AP-1 signaling pathways, including IL1 (IL1A and IL1B) and IL1 receptors (IL1R1, IL1R2, IL1RAP, and IL1RL1), IL6 (IL6 and LIF) and IL6 receptor, TNF and TNF receptors (TNFRSF10D, TNFRSF11B, and TNFRSF21), JAK2 and MAP kinases (MAP3K8 and MAP4K4) (Figure 3A). These genes are all common targets of STAT3, NFKB1, and AP-1 factors, with increased factor binding in promoters/enhancers (Figure 3B). Thus, the essence of the loop is that STAT3, NFKB1, and AP-1 directly activate upstream regulators that trigger the activation of intra-cellular signaling cascades, phosphorylate the transcription factors, and promote their nuclear localization and transcriptional activation. In accord with previous studies on IL6 (Iliopoulos et al., 2009), reducing the activation levels of JAK2, JNK, and IL1 receptor via inhibitors or antagonists results in decreased transformation efficiency (Figure S6). We define the 27 genes in the IL6/STAT3, IL1/NF-κB and TNF/AP-1 signaling pathways (Figure 3A) as the core positive feedback loop that maintains the transformed state.

**Figure 3.**
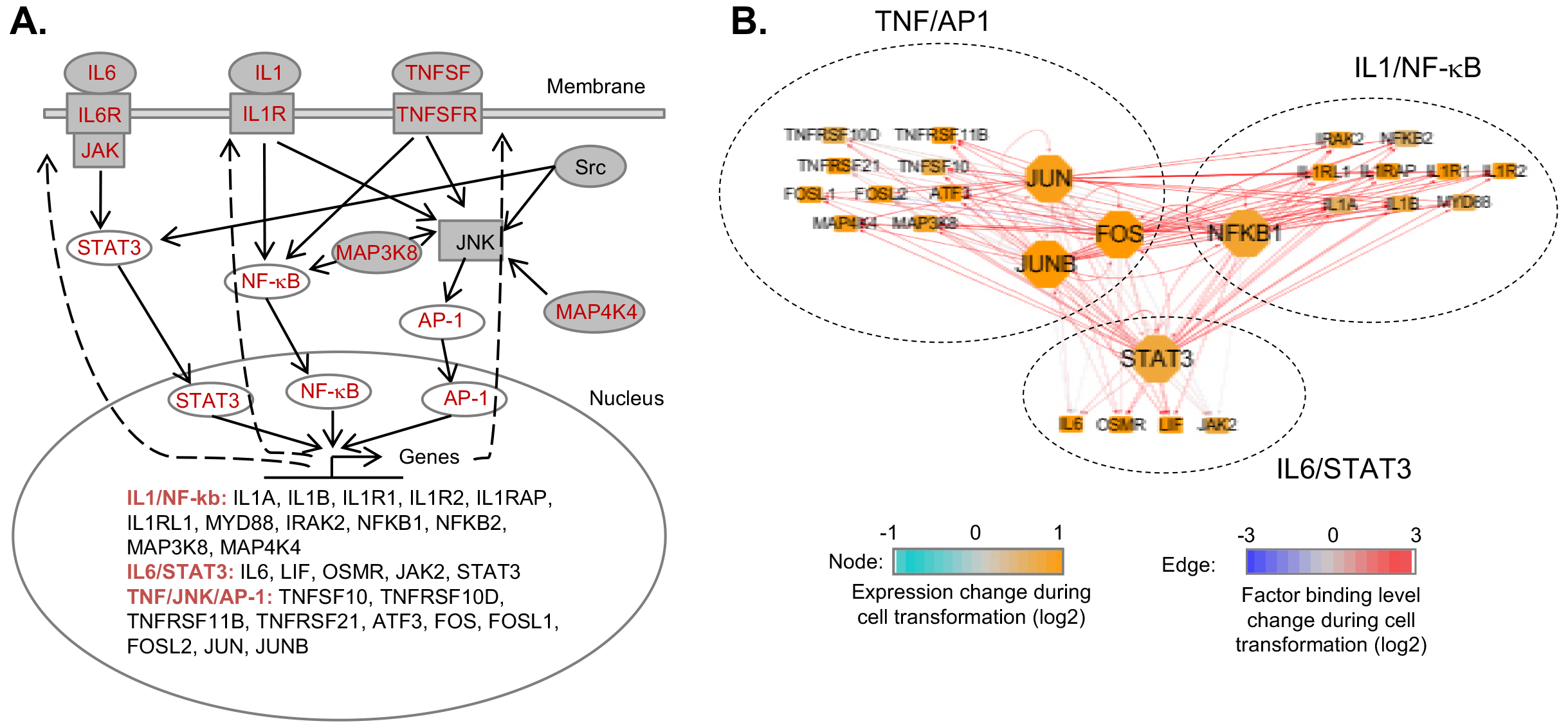
Positive feedback loop among IL1/NF-κb, IL6/STAT3 and TNF/AP-1 pathways. (A) The positive feedback loop of IL1/NF-κB, IL6/STAT3 and TNF/AP-1 pathways. The red colored genes are transcriptionally activated during transformation. (B) The network showing genes in the loop are common targets STAT3, NFKM, JUN, JUNB and FOS. The nodes represent genes in the loop. The edges represent binding of transcription factors in promoters/enhancers of target genes.

### An “inflammation index” to measure the inflammation level of a cancer cell line

The Cancer Cell Line Encyclopedia (CCLE) database contains gene expression data for 1,036 human cancer cell lines from over 20 developmental lineages (Barretina et al., 2012). The expression levels of 27 genes in the IL6/STAT3, IL1/NF-κB and JNK/AP-1 pathways (Figure 3) across those cancer cell lines (Figure 4A, B) are positively correlated beyond chance expectation (Median of Spearman’s Rank Correlation Coefficient = 0.17; Wilcoxon Rank Sum Test *P*-value < 10^−200^) (Figure 4C), indicating the inflammatory loop is relevant in different types of cancers. As expected from the extraordinary diversity of cancers, some cancer cell lines have transcriptional profiles much more similar to those our ER-Src model than others.

**Figure 4.**
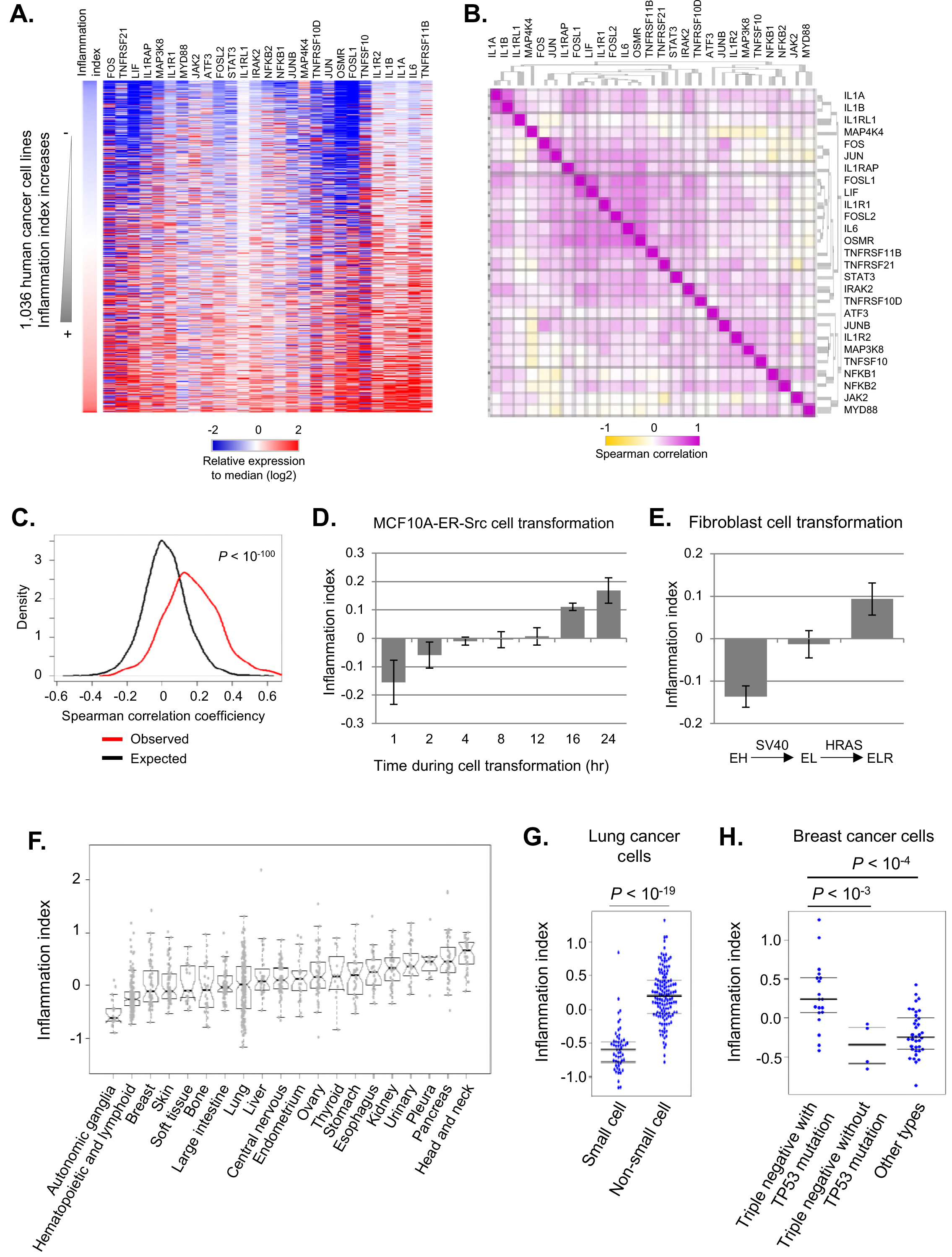
The inflammatory loop is active in different types of cancer cells. (A) Heatmap showing relative expression levels (median normalized values) of genes in the loop among 1,036 cancer cell lines. The samples were sorted by the inflammation index, which is calculated as the median value of the relative expression levels of the indicated 27 genes in the inflammatory loop. (B) Spearman correlation of expression levels of the gene pairs. (C) Distribution of observed and expected Spearman correlation coefficiencies of expression levels of genes in the loop. The expected correlation values were calculated by the randomly picked genes expressed in the cancer cell lines. (D) Change of inflammation index values during ER-Src cell transformation. Error bars represent standard deviation between two biological replicates. (E) Change of inflammation index values during fibroblast transformation. Error bars represent standard deviation between three biological replicates. (F) Distribution of inflammation index values among cancer cell lines from different developmental lineages. (G) Inflammatory indexes in genetic subtypes of lung cancer cell lines. (H) Inflammatory indexes in genetic subtypes of breast cancer cell lines.

To measure the inflammation level for each cancer cell line, we developed a scoring system called the “inflammation index”, which is calculated as the expression values of 27 genes (normalized to the median expression levels in the 1,036 cell lines) in the positive feedback loop. Importantly, this index is not simply based on a subset of genes that are induced during transformation, but rather genes that are direct targets of the STAT3/NF-κB/AP-1 regulatory network. The inflammatory levels gradually increase during ER-Src transformation (Figure 4D), and they are highly variable (>5 fold) among different cancer cell lines (Figure 4A). Similarly, in a fibroblast cell transformation model involving stepwise addition of the SV40 and RAS oncogenes to immortalized fibroblast, the inflammatory levels increase upon transformation (Figure 4E).

On average, head and neck as well as pancreatic cancer cell lines are most inflammatory, while the autonomic ganglia and blood cancer cell lines are least inflammatory (Figure 4F). However, cancer cell lines from the same developmental lineage can show high variance in inflammatory levels, which is correlated with their genetic subtypes. For example, cell lines from non-small cell lung cancers are more inflammatory than those of small cell lung cancers (Figure 4G), and triple negative breast cancer cell lines with p53 mutations are more inflammatory than other subtypes (Figure 4H).

### The inflammatory loop is active in tumors from cancer patients

Although cancer cell lines are derived from tumors, long-term propagation of cell lines under artificial conditions raises the possibility that cell line data might be misleading with respect to cancer. We addressed the relevance of the inflammatory loop in human tumors using RNA-seq data in the Cancer Genome Atlas database (Cancer Genome Atlas Research et al., 2013). In accord with the results in cancer cell lines, genes in the IL6/STAT3, IL1/NF-κB and JNK/AP-1 inflammatory loop are co-expressed in human breast tumors (Figure 5A). Similarly, triple negative breast tumors are more inflammatory than other types of breast tumors (Figure 5B). Moreover, the median inflammatory index value of all tumor samples from a given developmental lineage is highly correlated with that of cells lines from the same developmental lineage (Pearson correlation coefficient = 0.82; *P*-value < 10^−5^) (Figure 5C). As we combined all tumors for a given lineage, it seems unlikely that this analysis is significantly compounded by differences in the tumor microenvironment. These observations indicate that, with respect to inflammation and gene regulation profiles, cancer cell lines are good models for tumors, and the inflammatory loop is very relevant for many types of human cancer.

**Figure 5.**
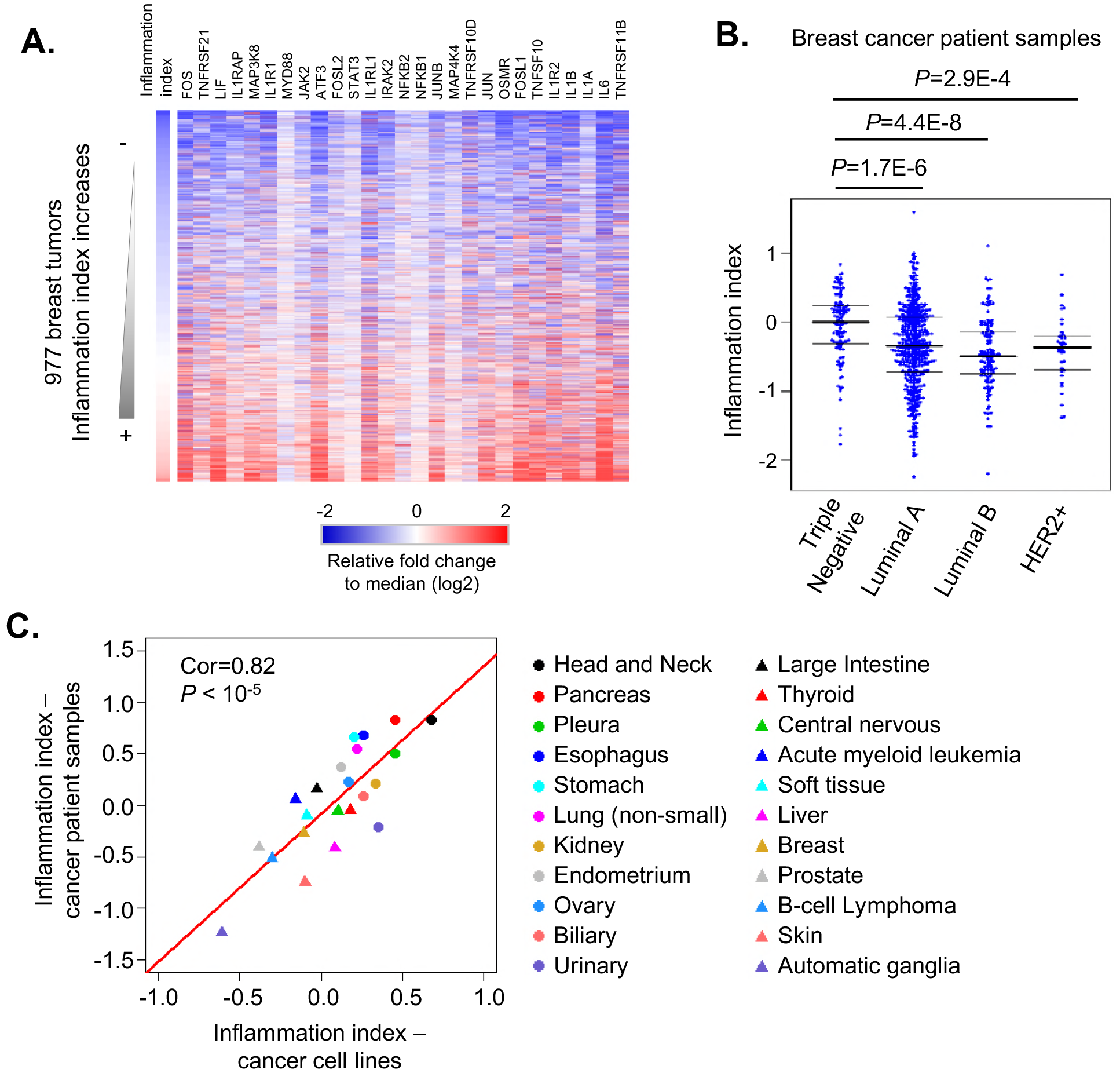
The inflammatory levels are variable among cancers with different developmental lineages and genetic subtypes. (A) Heatmap showing the relative expression levels of 27 genes in IL6/STAT3, IL1/NF-κB and TNF/NF-κB pathways in breast cancer patient samples from the TCGA database. (B) Inflammatory indexes in genetic subtypes of breast cancer patients. (C) Correlation between inflammatory levels between cancer cell lines and patient tissue samples among different developmental lineages. The median inflammation index value of cancer samples from the same developmental lineage indicates its inflammatory level.

### Identification of other genes whose expression is correlated to the inflammation index

Previous analyses of the ER-Src model designed to identify oncogenically relevant genes have relied on differential gene expression upon transformation and/or direct regulation by NF-κB, STAT3, and AP-1 factors. However, it is highly likely that this approach will miss critical genes that are part of the tumor-promoting inflammation process. As an alternative approach, for every gene, we calculated the Spearman correlation coefficient between its expression level and the inflammation index across cancer cell lines for various developmental lineages. The correlation values are significantly conserved among developmentally distinct cancers, again indicating a widespread role of the inflammatory loop and it target genes (Figure 6A).

**Figure 6.**
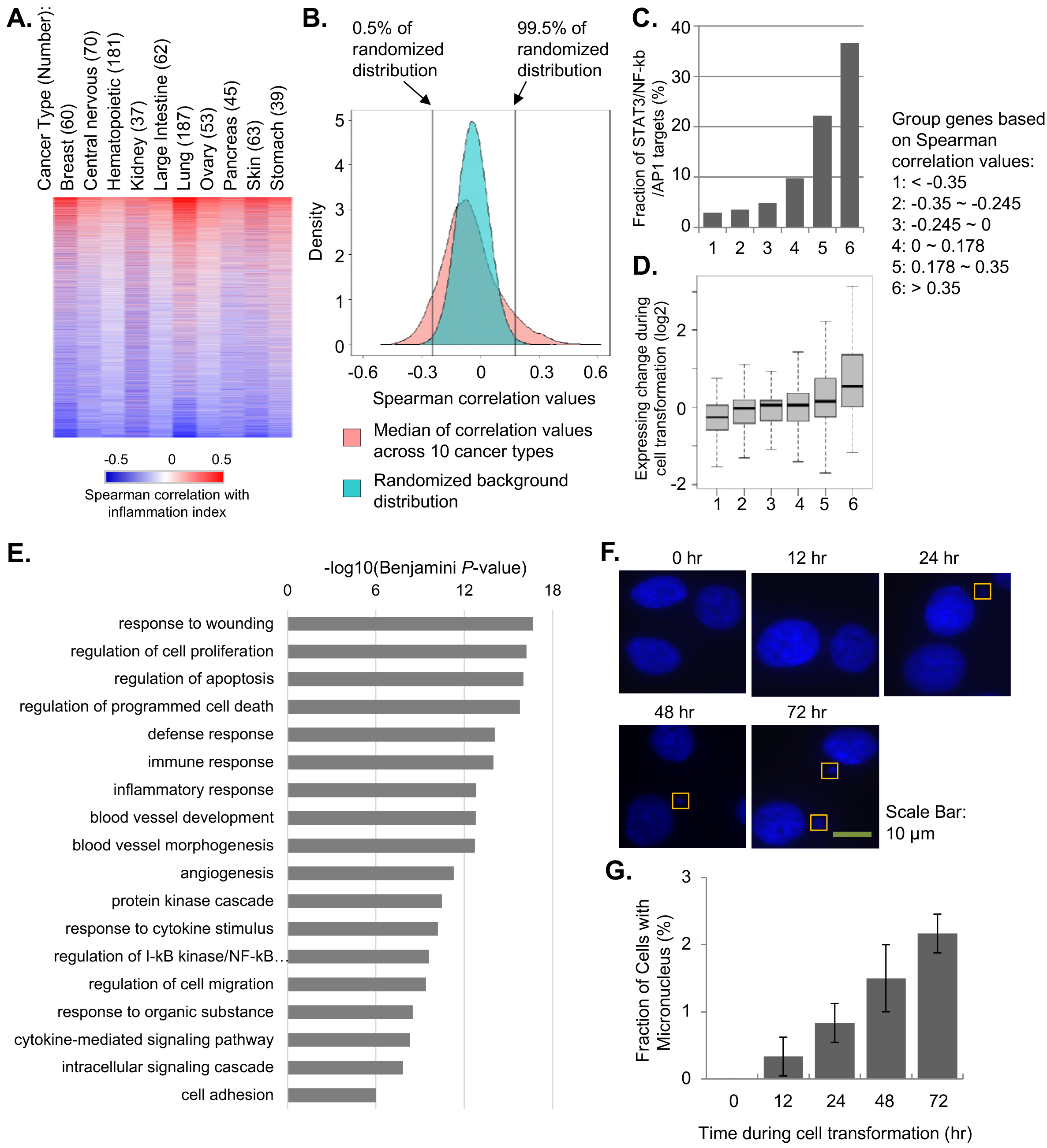
Genes and pathways with expression levels correlated with inflammatory loop. (A) Heatmap showing the Spearman correlation coefficiencies between gene expression levels and inflammatory index across 10 types of cancers. (B) Distribution of the median correlation values across 10 cancer types. For the background distribution, we randomized the correlation values in each cancer type, and calculated the median values of the randomized values. (C) Fractions of genes are direct targets of STAT3/NF-κB/AP1. Genes were grouped based on Spearman correlation values with inflammation index as in (B). (D) Expression change of the gene groups during transformation. (E) Gene ontology of genes showing significant positive correlation of expression and inflammatory index (median Spearman correlation coefficiencies across cancer types > 0.18). (F) Fluorescent nuclear imaging of Micronuclei reveals increased genome instability. (G) Fraction of cells showing micronucleus formation during transformation.

Genes showing higher positive correlation with the inflammation index are more likely to be direct transcriptional targets of STAT3/NF-κB/AP1 and tend to be upregulated during transformation (Figure 6B, 6C and 6D). We identified 1,303 genes showing significant positive correlation of expression with the inflammation index (median Spearman’s Rank Correlation Coefficient across cancer cell types > 0.18, false discovery rate < 0.005)(Figure 6B). Aside from genes involved in the inflammatory response, these genes are enriched in biological pathways such as angiogenesis, cell proliferation, apoptosis, intracellular signaling cascade and cell migration (Figure 6E). These pathways, which are defined by the inflammatory index and not transformation *per se*, are in excellent accord with the pathways accord with pathways activated during ER-Src transformation (Figure 6E), thereby providing independent evidence that the inflammatory loop and oncogenic pathways uncovered in the ER-Src cell transformation model are highly relevant for different types of human cancer.

### A correlation between inflammation and genome instability in cancer cell lines

1,369 genes show negative correlation between expression levels and inflammation index, and these are enriched in pathways including DNA metabolic process, DNA replication, DNA repair and cell cycle (Benjamin FDR < 10^−13^). This observation suggests that oncogenic-associated inflammation is inversely related to genome stability, and indeed the genes showing most negative correlation (e.g. MSH2, FANCF, BRCA1) are regulators of genome instability. Interestingly, the genes related to genome instability are not transcriptionally regulated during ER-Src cell transformation.

To examine the link between inflammation and genome instability, we performed micronucleus staining of cells during ER-Src transformation. Indeed, more cells contain micronuclei as transformation processes (Figure 6F, G), with 2.2% cells containing micronuclei after 72 hr after induction of transformation as compared to virtually no cells before transformation. Thus, the inflammation-mediated process of transformation is associated with decreased genome stability.

### Inflammatory tumor samples contain increased levels of non-cancer cells

As shown above, the inflammation indices of cancer cell lines for a particular cancer type are highly correlated with the corresponding tumor samples (Figure 5C). However, unlike samples from cancer cell lines, tumor samples not only contain the cancer cells, but also immune and stromal cells from the tumor microenvironment. Such tumor impurity complicates the analysis of gene expression profiles of tumor cells, particularly as immune cells are also inflammatory. We therefore took several approaches to disentangle the inflammatory nature of cancer cells from other cells in a tumor.

First, for numerous tumor samples we analyzed the relationship between the inflammatory index and tumor purity (fraction of cancer cells in a sample), which has been estimated from cancer-specific genetic mutations or cell type-specific gene signatures (Aran et al., 2015). In general, the inflammation level increases as the tumor purity decreases (Figures 7A and S7A-D). However, triple negative breast tumors have higher inflammatory levels than those of other genetic subtypes, even though all of these forms of breast cancer have similar purity levels (Figure S7E). The higher inflammatory level of triple negative breast tumors is consistent with the results in the corresponding cancer cell lines (Figure 5B), indicating that it reflects the intrinsic properties of the cancer cells and not the degree of tumor purity.

**Figure 7.**
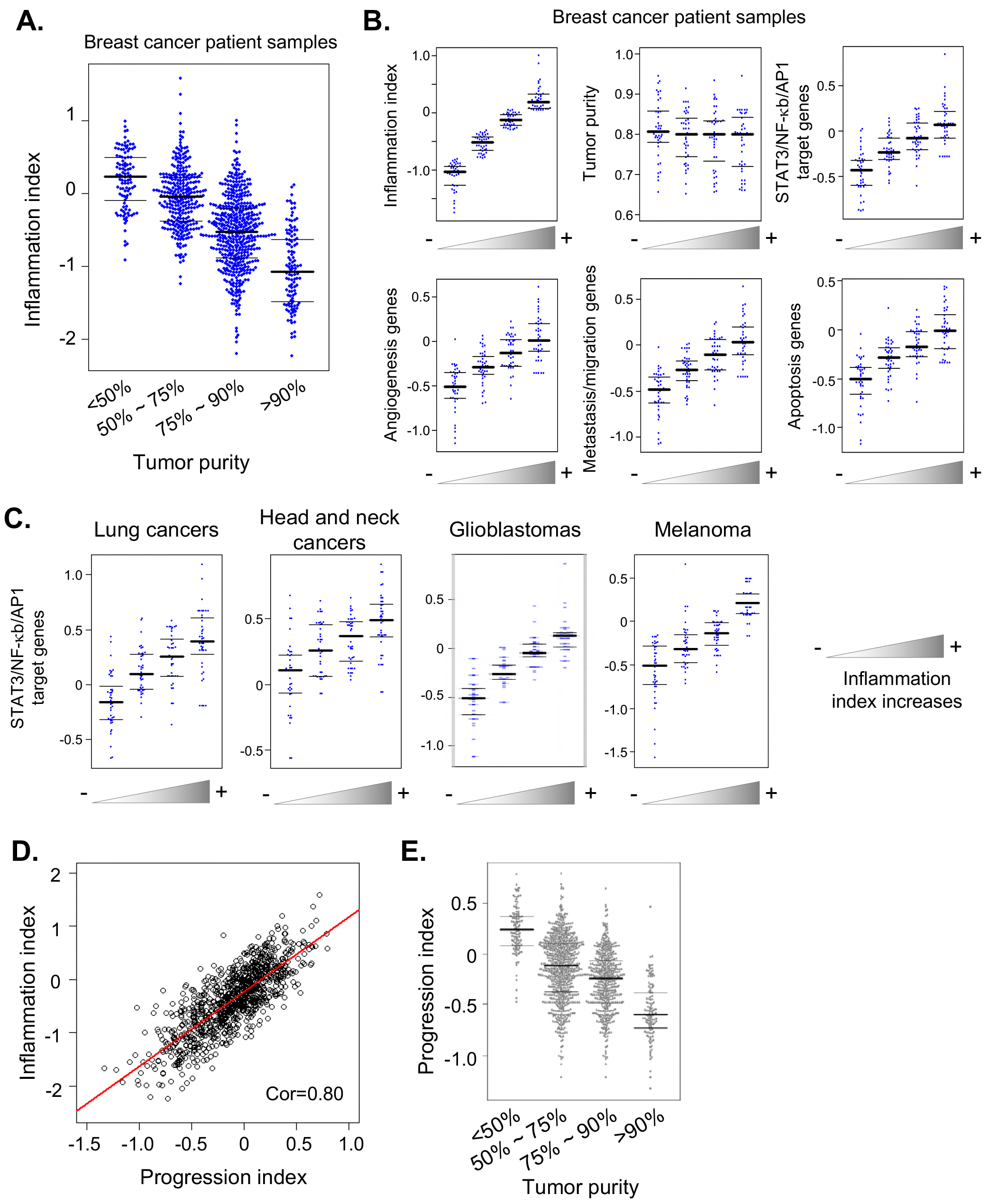
Correlation between inflammation index, cancer progression index and tumor purity. (A) Correlation between tumor purity and inflammation index of breast cancer patient samples. (B) Randomly picked breast tumor samples with different inflammation levels and similar purity were examined for gene expression levels in the indicated oncogenic pathways that are direct targets of STAT3/NF-κB/AP-1 and also show significant positive correlation of expression with inflammation index in Figure 6B. Genes associated with immune/inflammatory response were removed from the analyses. (C) Randomly picked tumor samples with different inflammation levels and similar purity were examined for expression levels of target genes of STAT3/NF-κB/AP1 in lung cancers, head and neck cancers, glioblastomas and melanoma. (D) Correlation between inflammation index and cancer progression index. (E)Correlation between tumor purity and cancer progression index.

As an alternative approach, we selected breast patient samples with similar levels of tumor purity but different levels of inflammation (divided into 4 bins) and analyzed expression of non-inflammatory genes whose expression is strongly correlated with the inflammatory index (Figure 7B). Expression of these transformation-related, but noninflammatory genes, which are direct targets of STAT3/NF-κB/AP1 and are involved in angiogenesis, apoptosis and cell migration, increases as the inflammation index increases. Similar results are observed in all other types of cancer we examined such as melanoma, lung, head and neck, and glioblastomas (Figures 7C and S8). These results indicate that the inflammatory loop and associated network are active in various types of cancers, although we can’t exclude the formal possibility that non-malignant cells in the various samples might be in different states.

Lastly, we created a “progression index” that is strongly correlated with the inflammatory index (Figure 7D), but is based on 98 genes that regulate “migration/metastasis”, “apoptosis” and “angiogenesis”, but are not currently annotated as inflammatory or part of the immune response. This progression index, should reflect the regulatory circuits involved in cellular transformation and tumor formation, but not normal immune cells. Indeed, across the large set of tumor samples, the score of non-inflammatory index inversely correlated with tumor purity (Figure 7E). All of these observations indicate that tumors containing inflammatory cancer cells preferentially contain non-cancer cells. Thus, tumor-associated inflammation level is positively correlated with the complexity of microenvironment and the presence of immune cells.

## DISCUSSION

### NF-κB, STAT3, and AP-1 factors are the core of the positive feedback loop controlling tumor-promoting inflammation

In the ER-Src cellular transformation model, a transient inflammatory stimulus mediates an epigenetic switch from a stable non-transformed cell to a stable transformed cell (Iliopoulos et al., 2009). Epigenetic switches are the basis of multicellular development, and they occur by activating a positive feedback loop that maintains the altered state. In the ER-Src model, Src activates the inflammatory transcription factors STAT3 and NF-κB that form the basis of the inflammatory feedback loop that is required for maintenance of the transformed state (Iliopoulos et al., 2009; Iliopoulos et al., 2010; Fleming et al., 2015; Ji et al., 2018).

Here, we show that AP-1 factors play a critical role in the inflammatory feedback loop. AP-1 factors are not only important for transformation, but they form complexes with STAT3 and/or NF-κB that bind target sites. Specifically, these factors co-immunoprecipitate, and their binding profiles are coincident at many target sites. Most, and perhaps all, of the sites where co-binding occurs contain AP-1 motifs, suggesting that the AP-1 factors directly interact with DNA, whereas STAT3 and NF-κB are often recruited via interactions with the AP-1 factors. At some sites, STAT3 and NF-κB bind via their own motifs in the absence of AP-1 factors. At present, it is unclear whether AP-1, NF-κB, and STAT3 can form a ternary complex at individual sites or if NF-κB and STAT3 form independent complexes with AP-1 factors at these sites.

In addition to their roles in inflammation *per se*, STAT3, NF-κB and AP-1 work together to regulate key genes in oncogenic pathways such as angiogenesis, apoptosis, cell migration and epithelial to mesenchymal transition. Expression of many genes in these pathways is induced upon transformation in a manner that is linked to increased binding of all these factors. Moreover, expression of many common genes is reduced upon knockouts of individual factors, indicating that all of these factors contribute to expression of these genes. Our identification of a common STAT3/NF-κB/AP-1 network is distinct from, but not inconsistent with, previous observations that STAT3 and NF-κB have different (i.e. non-overlapping) binding sites and gene expression effects during the transformation process (Fleming et al., 2015).

The positive feedback loop elucidated here is considerably more complex than previously described, and it is maintained in two distinct ways. The key transcription factors directly bind and activate the expression of each other, thereby reinforcing the core transcriptional state. This mutual co-regulation of key transcription factors is similar to what occurs in muscle development (MyoD and related factors), embryonic stem cells (Nanog, Oct4), and presumably all epigenetic states. In addition, these transcription factors directly regulate nearly all upstream components in the IL6/STAT3, IL1/NF-κB and TNF/AP-1 signaling pathways, thereby maintaining the activity of the individual factors and hence the gene regulatory pattern. Thus, this extensive positive feedback loop represents a coherent regulatory system in which numerous interconnected components stably maintain a common, yet complex, transcriptional program

### The inflammatory gene signature as an approach to type human cancers

Historically, cancer types were classified by their developmental origin as well as by crude cellular phenotypes. More recently, cancers have been classified by the genetic mutations that drive the oncogenic state. Such classification, together with drugs targeted to specific mutations, has been the primary basis for the idea of personalized medicine approaches to cancer treatment. Here, we develop a functional approach to classify human cancers that utilizes gene expression signatures, specifically an inflammation index based on the STAT3/NF-κB/AP-1 network. Importantly, this index is based on an integrated regulatory network and hence is significantly different from indices based solely on transcriptional profiles that arise from multiple regulatory inputs.

Overall, expression levels of the 27 key genes in the IL6/STAT3, IL1/NF-κB and TNF/AP-1 transcription loop that make up the inflammation index are strongly correlated with each other. Not only does this observation provide further support for the coherence of the feedback loop, but it strongly suggests that the loop and associated network functions to various degrees in different cancer cell lines and tumors. Transformed cell lines have a higher inflammation index than observed with non-transformed cells, suggesting a general role of inflammation in cancer. However, there is a wide range of inflammation scores among various cancers that occurs in developmentally unrelated cancers and is not strictly linked to specific mutations. Importantly, the inflammation indices of cell lines and tumors are strongly related, indicating its relevance for human disease. We predict the existence of drugs whose efficacy correlates with the inflammation index (and by extension other gene expression indices). If so, the inflammation index of the tumor cells might be useful for choosing drugs for personalized cancer treatment.

### Inflammatory cancers may recruit immune cells to the tumor site

Patient tumor samples contain cancer cells and immune (and other non-cancer) cells, all of which contribute to the transcriptional profile. Interestingly, over a large number of tumor samples, there is an inverse relationship between the inflammatory index and the estimated degree of sample purity. In principle, this relationship could merely reflect the fact that immune cells also express many inflammatory genes, which could significantly contribute to the observed inflammation index of the sample.

We attempted to distinguish the contributions of cancer and immune cells to the transcriptional profile by analyzing many non-inflammatory genes (either in specific pathways or as an overall index) whose expression is very strongly correlated with the inflammatory index and hence to the STAT3, NF-κB, AP-1 regulatory network. These observations suggest a dynamic interplay between cancer cells and immune cells that is linked to the inflammation index of the cancer cells in the sample. We propose that cytokines and chemokines secreted by the cancer cells attract immune cells to their vicinity of the tumor, thereby resulting in a less pure tumor sample. In addition, these cytokines and chemokines secreted by the cancer cells also help activate the inflammatory gene expression program in immune cells. Thus, we propose that the mutual and perhaps synergistic functional interactions between cancer cells and immune cells can create an inflammatory microenvironment in a manner that depends on the inflammatory properties of the cancer cells.

## MATERIALS AND METHODS

### Cell culture

The inducible model of cellular transformation involves MCF-10A, a non-transformed mammary epithelial cell line (Soule et al., 1990) containing ER-Src, a derivative of the Src kinase oncoprotein (v-Src) that is fused to the ligand-binding domain of the estrogen receptor (Aziz et al., 1999). Cells were cultured in DMEM/F12 medium with the supplements as previously described (Iliopoulos et al., 2009; Hirsch et al., 2010). Tamoxifen (Sigma, H7904, 0.4 mM) was used to transform this inducible cell line, when the cells were grown to 30% confluence. For the drug inhibition experiments, cells were pretreated with SP600125 (JNK inhibitor, Selleck Chemical, S1460, 0.3 μM), S-Ruxolitinib (JAK inhibitor, Cayman Chemical, #11609, 0.1 μM), Tofacitinib (JAK inhibitor, Selleck Chemical, S2789, 0.1 μM) and IL-1RA (IL-1R antagonist, Peprotech, #200-01RA, 30 ng/ml) for 75 minutes after which cells were transformed by addition of 0.4 mM Tamoxifen and 4 ng AZD0530 for 24 hours. Transformation efficiency was measured by cell growth under low attachment conditions as described previously (Rotem et al., 2015).

### CRISPR Knockouts

CRISPR-blasticidin lentiviral plasmid was constructed by replacing puromycin resistance gene with blasticidin resistance gene in LentiCRISPR V2 plasmid (Addgene, #52961). Table S2 is the list of the oligo sequences used to clone into CRISPR-blasticidin plasmid. CRISPR-blasticidin plasmid and three lentiviral plasmids, VSV-G, GP and REV were co-transfected into 293T cells to produce lentiviruses as the previous publication (He et al., 2011). After CRISPR lentiviruses infection, ER-Src cells were selected with blasticidin 10 μg/ml for 3 days to generate CRISPR knockout stable cell lines. The knockout efficiencies of the transcription factors were assessed by Western blotting.

### Nucleocytoplasmic separation, co-immunoprecipitation, and Western blotting

Cells were suspended in a buffer (10 mM HEPES PH 7.5, 10 mM KCl, 2 mM MgCl, PMSF 0.1 mM and Roche complete protease inhibitor, Sigma, #11697498001) and incubated on ice for 20 min. After grinding the cells 50 times with a Wheaton Dounce A, the cell lysate was layered on top of 40% sucrose buffer and centrifuged at top speed at 4°C for 5 minutes. The supernatant (cytoplasm) and pellet (nucleus) were separated, and the The nuclear pellet was re-suspended in 20 mM Tris pH 7.4, 150 mM NaCl, 1 mM EDTA, 1 mM EGTA, 1% Triton X-100, 25 mM sodium pyrophosphate, 1 mM NaF, 1 mM β-glycerophosphate, 0.1 mM sodium orthovanadate, 1 mM PMSF, 2 μg ml-1 leupeptin and 10 μg ml-1 aprotinin. Co-immunoprecipitations were performed by mixing nuclear and cytoplasmic fractions with antibodies and Dynabeads protein G (Life Technology, 10004D) in buffer (50 mM Tris (PH 7.5), 100 mM NaCl, 1.5 mM EGTA and 0.1% Triton X-100) at 4°C overnight. Dynabeads were washed with Co-IP buffer for 8 times, and immunoprecipitated proteins were analyzed by Western blotting. Antibodies for co-immunoprecipitations were against FOS (Cell Signaling, #2250), NF-κB1 (SCBT, SC-372X), and STAT3 (Cell Signaling, #9139), and antibodies for western blotting were against STAT3 (Cell Signaling, #12640), RELA (Cell Signaling, #8242), and JUNB (Cell Signaling, #3753 and SCBT, SC-8051).

### Micronucleus assays

Cells were seeded into 8-well chamber slides (LAB-TEK, #154941), transformed for indicated times, fixed with 4% Formaldehyde, and then stained with DAPI (2 μg/ml) for 5 minutes. The pictures were captured using Michael Widefield inverted Nikon Ti2 fluorescence microscope at Nikon Imaging Center, Harvard Medical School. Micronuclei were counted from 5 random fields of each time point.

### ChIP-seq and DNase-seq to define chromatin states

ChIP-seq (Fleming et al., 2015) and DNsae-seq (Thurman et al., 2012) were performed as described previously. Fastq reads were aligned to human reference genome (hg19) using Bowtie (Langmead et al., 2009) allowing up to 2 mismatches. Only the uniquely mappable reads were used for subsequent analyses. For ChIP-seq data for STAT3, NFKB1, JUN, JUNB, FOS, H3K27ac, H3K4me3 and H3K4me1, we used MACS (Zhang et al., 2008) to call peaks with the cutoff *P*-value < 10^−8^ in at least one sample, using the parameters “macs2 callpeak --llocal 1000000 -g 2.7e9”. For ChIP-seq data for H3K27me3, H3K9me3 and H3K36me3, we used SICER (Zang et al., 2009) to call peaks with the cutoff E-value > 40, window size 200 bp and gap size 600 bp, which is better for identifying broad read peaks. For DNase-seq data, we used MACS (Zhang et al., 2008) to call peaks with the cutoff *P*-value < 10^−11^ in at least one sample and using the following parameters “macs2 callpeak --llocal 1000000 -g 2.7e9”. Chromatin states were defined as promoters (H3K4me3 peaks), active enhancers (H3K27ac peaks, but no H3K4me3), poised enhancers (H3K4me1 peaks, but no H3K27ac or H3K4me3 peaks) or heterochromatin (H3K9me3 or H3K27me3 peaks).

### Analyses of transcription factor binding

Transcription factors tend to localize into *cis*-regulatory regions (CRRs) that regulate gene expression. We merged overlapping peaks of all factors to define CRR regions. For each CRR, we measured factor binding levels as Reads per Million (RPM) using ChIP-seq data, and chromatin accessibility based on DNase-seq data. The peak summit of each factor binding site was defined based on MACS (Zhang et al., 2008). In Figure 1B, we plotted the distance between peak summits of paired transcription factors, located in the same CRR. For each factor binding site, we took 50 nt around the peak summits to perform the motif analyses, using the HOMER (Heinz et al., 2010). We used the position weight matrices (PWM) of STAT, NF-κB and AP1 in HOMER (Heinz et al., 2010). In Figure 1D, we plotted the distribution of STAT, NF-κB and AP1 motifs around peak summits of transcription factors. As the control, we shuffled the nucleotide positions of PWM, and kept the A/T/G/C occurrence frequency of the motifs as the same. We created the shuffled motifs for 50 times for each motif and plotted their occurrence around the factor peak summits. To examine the contribution of factor binding to gene expression, we assigned factor binding peaks to the closest expressed genes within a distance of 200 megabases, summed up the ChIP signal, and calculated the binding level fold-change during transformation.

### Transcriptional profiling

RNA was extracted using mRNeasy Mini Kit following the manufacturer’s instruction. RNA-seq libraries were prepared using TruSeq Ribo Profile Mammalian Kit (Illumina, RPHMR12126) as per manufacturer’s instruction. RNA-seq libraries were sequenced by Harvard Bauer Core Facility using Hiseq 2000. Raw reads were aligned to GENCODE (Harrow et al., 2012) defined transcripts and then human reference genome (hg19) using Tophat (Langmead et al., 2009) allowing up to 2 mismatches. The gene expression levels were indicated as transcript per million (TPM) values. For Figure 4D and 4E, we downloaded microarray data to from GSE17941 to examine gene expression dynamics during MCF10-ER-Src and fibroblast cell transformation (Hirsch et al., 2010). The gene expression values were calculated by the RMA approach using Affymetrix Expression Console Software.

### CCLE and TCGA data analyses

The gene expression, genetic and lineage annotations of 1,036 cancer cell lines were downloaded from the Cancer Cell Line Encyclopedia (CCLE) (Barretina et al., 2012). The level 3 data showing clinical annotations and RNA expression of cancer patients were downloaded from TCGA database. The tumor purity estimations were obtained from (Aran et al., 2015), which estimated tumor purity using 5 different measurements: STIMATE, based on gene expression profiles of 141 immune genes and 141 stromal genes; ABSOLUTE, based on somatic copy-number data; LUMP, based on 44 non-methylated immune-specific CpG sites; IHC, based on image analysis of haematoxylin and eosin stain slides produced by the Nationwide Children’s Hospital Biospecimen Core Resource; and the averaged values based on 4 methods above.

**Calculation of inflammation index.** The 27 signature genes in the inflammatory loop include genes in IL1/NF-κB pathway (IL1A, IL1B, IL1R1, IL1R2, IL1RAP, IL1RL1, MYD88, IRAK2, NFKB1 and NFKB2), IL6/STAT3 pathways (IL6, LIF, OSMR, JAK2 and STAT3), TNF /AP-1 pathway (TNFSF10, TNFRSF10D, TNFRSF11B, TNFRSF21, ATF3, FOS, FOSL1, FOSL2, JUN and JUNB) and MAP kinases (MAP3K8 and MAP4K4). We calculated Spearman’s rank correlation coefficient values between gene pairs using gene expression data from CCLE and TCGA. For Figure 4C, we randomly selected expressed genes and calculated Spearman’s correlation as the background distribution. We grouped breast cancer patients based on their genetic subtypes as following: Triple Negative (ER-, PR-and HER2-), Luminal A (ER+, PR+ and HER2−), Luminal B (ER+, PR+ and HER2+), and HER2+ (ER−, PR− and HER2+). To calculate the inflammation index based on these genes, we first normalized the log2 expression levels across all samples to their median values. Then for each sample, we calculated the median expression level of signature genes representing the inflammation index.

### Calculation of cancer progression index

We picked 98 genes showing significant positive correlation with inflammation index across different cancer types (Figure 6B), are linked to “cell migration”, “angiogenesis” and “apoptosis”, and are not immune/inflammation related based on gene ontology definition. These genes include: ACTC1, ACTN4, ADAM10, ADAMTSL4, AHR, ANXA2, ARHGAP22, C8ORF4, CARD6, CAV1, CFLAR, CIB1, COL15A1, COL1A1, COL4A2, CSRNP1, CYP1B1, CYR61, DAB2, DAP, DFNA5, DLC1, DRAM1, ECE1, EDN1, ELK3, EPAS1, EPHB2, F3, FN1, GADD45A, GADD45B, HBEGF, HIPK3, HMOX1, HSPG2, HTATIP2, IER3, IGFBP3, ILK, ITGA5, ITGAV, JAG1, JUN, LAMB1, LAMC2, LEPR, LGALS1, MYADM, MYH9, MYO1C, NEK6, NFKBIA, NOTCH2, NRP1, NUMB, PEA15, PHLDA1, PHLDA2, PLAU, PLK3, PNPLA6, PPP1R13L, PPP1R15A, PRKCA, PTPRB, PTPRH, RALB, RHBDD1, RRAS2, RTN4, SDCBP, SEMA4B, SERPINB3, SGK1, SH3GLB1, SH3KBP1, SH3RF1, SHB, SHC1, SLC12A6, SQSTM1, SRPX2, STEAP3, STK17A, STK17B, STX4, TGFA, TGFBI, TGFBR2, TMEM214, TSPO, UBE2Z, VEGFA, WARS, WASF2, XAF1 and ZFP36L1. We used the same calculation steps described for the inflammation index to determine the cancer progression index.

### Gene Ontology analyses

The Database for Annotation, Visualization and Integrated Discovery (DAVID) (Huang da et al., 2009) was used for gene ontology analyses.

## Data availability

All sequencing data that support the findings of this study have been deposited in the National Cancer for Biotechnology Information Gene Expression Omnibus (GEO) and are accessible through the GEO series accession numbers GSE115597, GSE115598 cand GSE115599. All computational codes are available from the authors upon request.

## ACKNOWLEDGEMENTS

This work was supported by the Searle Leadership Fund in the Life Sciences from Northwestern University to Z.J., a Career Transition Award to Z.J. from the National Cancer Institute (K99 CA 207865), and a research grant to K.S. from the National Institutes of Health (CA 107486). A.R. is a Howard Hughes Investigator.

## SUPPLEMENTARY FIGURE LEGENDS AND TABLES

**Figure S1.**
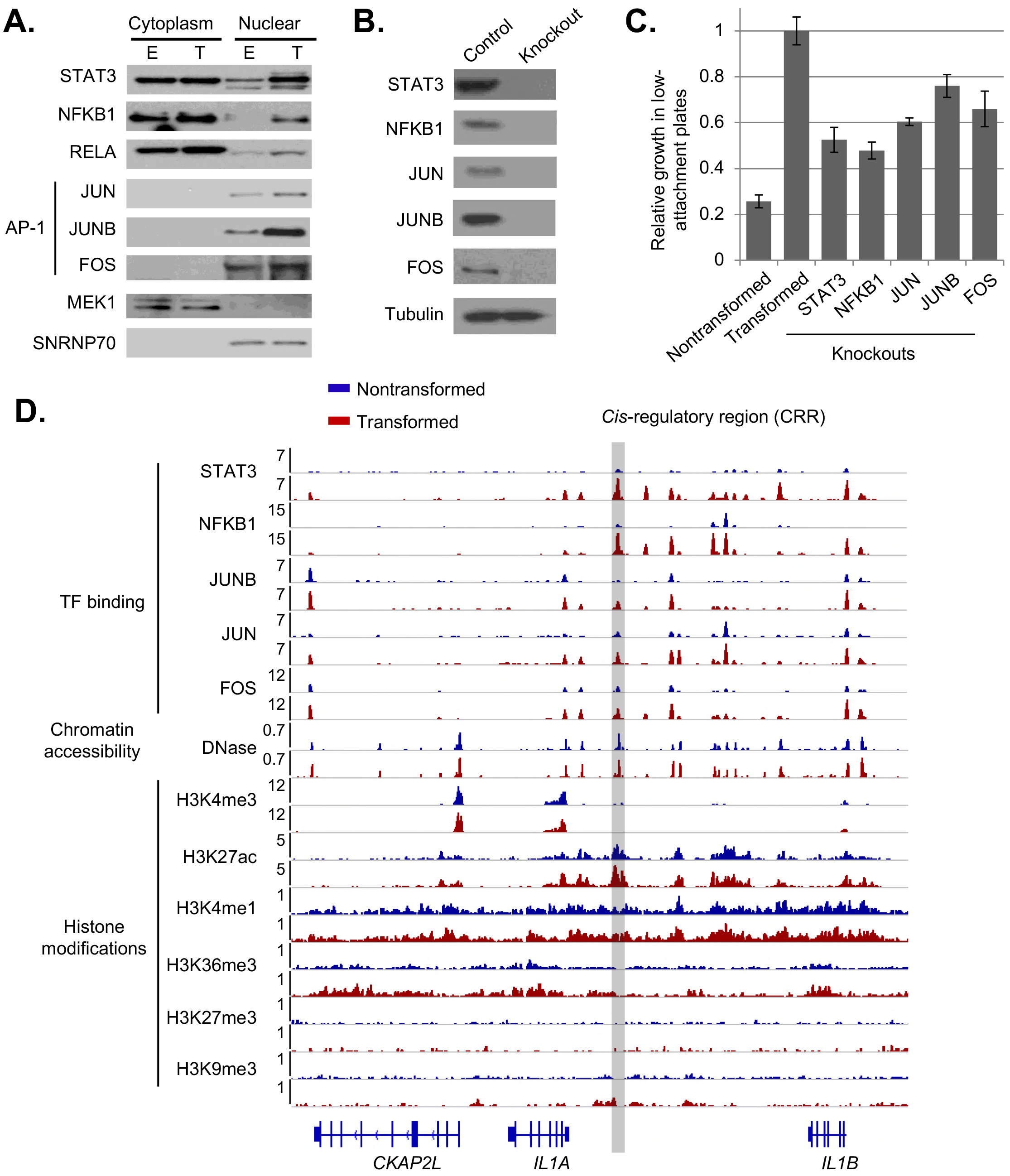
ChIP-seq for STAT3, NFKB1, JUN, JUNB and FOS during MCF10A-ER-Src cell transformation. (A) Western blots showing expression of the indicated proteins in the nucleus and cytoplasm during transformation. (B) Western blots confirming CRISPR knockouts of the indicated transcription factors. Tubulin was used for the control. (C) Cell transformation assays for cells upon factor knockouts and the control. (D) A genomic region showing ChIP-seq data for the indicated transcription factors and histone modifications and DNase-seq data for measuring chromatin accessibility before and after transformation.

**Figure S2.**
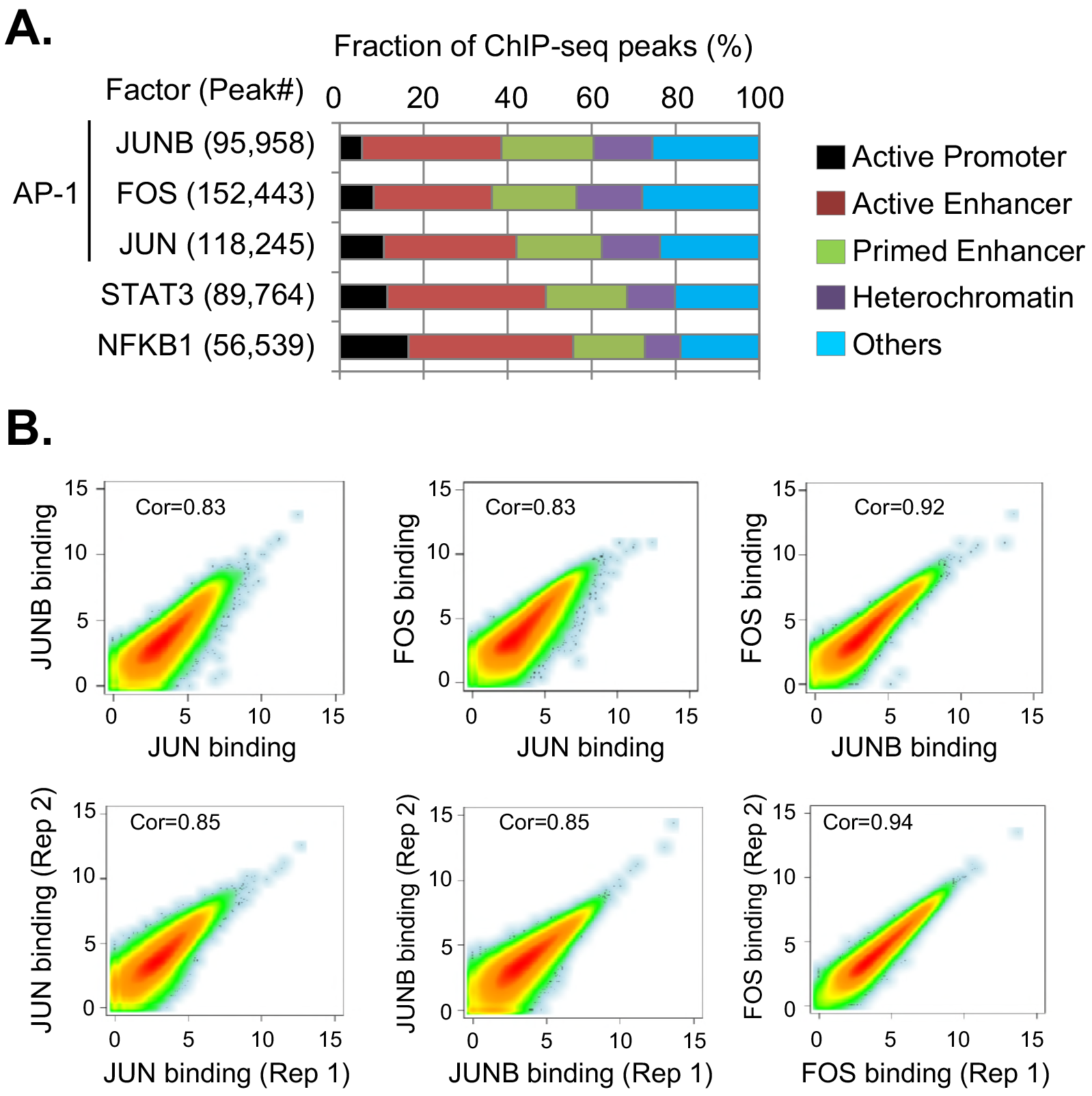
Genomic distribution of STAT3, NF-κB and AP-1 binding sites. (A) Genomic locations of binding sites for the indicated transcription factors. Regulatory regions were classified by chromatin status as follows: active promoters (H3K4me3); active enhancers (H3K27ac, but no H3K4me3); primed enhancer (H3K4me1 only); Heterochromatin (H3K9me3 or H3K27me3); others (without histone modification peaks). (B) Correlation of JUN, JUNB and FOS binding levels, and correlation of binding levels between biological replicates. We merged overlapping peaks for these AP-1 factors for the analyses. The Pearson correlation coefficiency values are shown.

**Figure S3.**
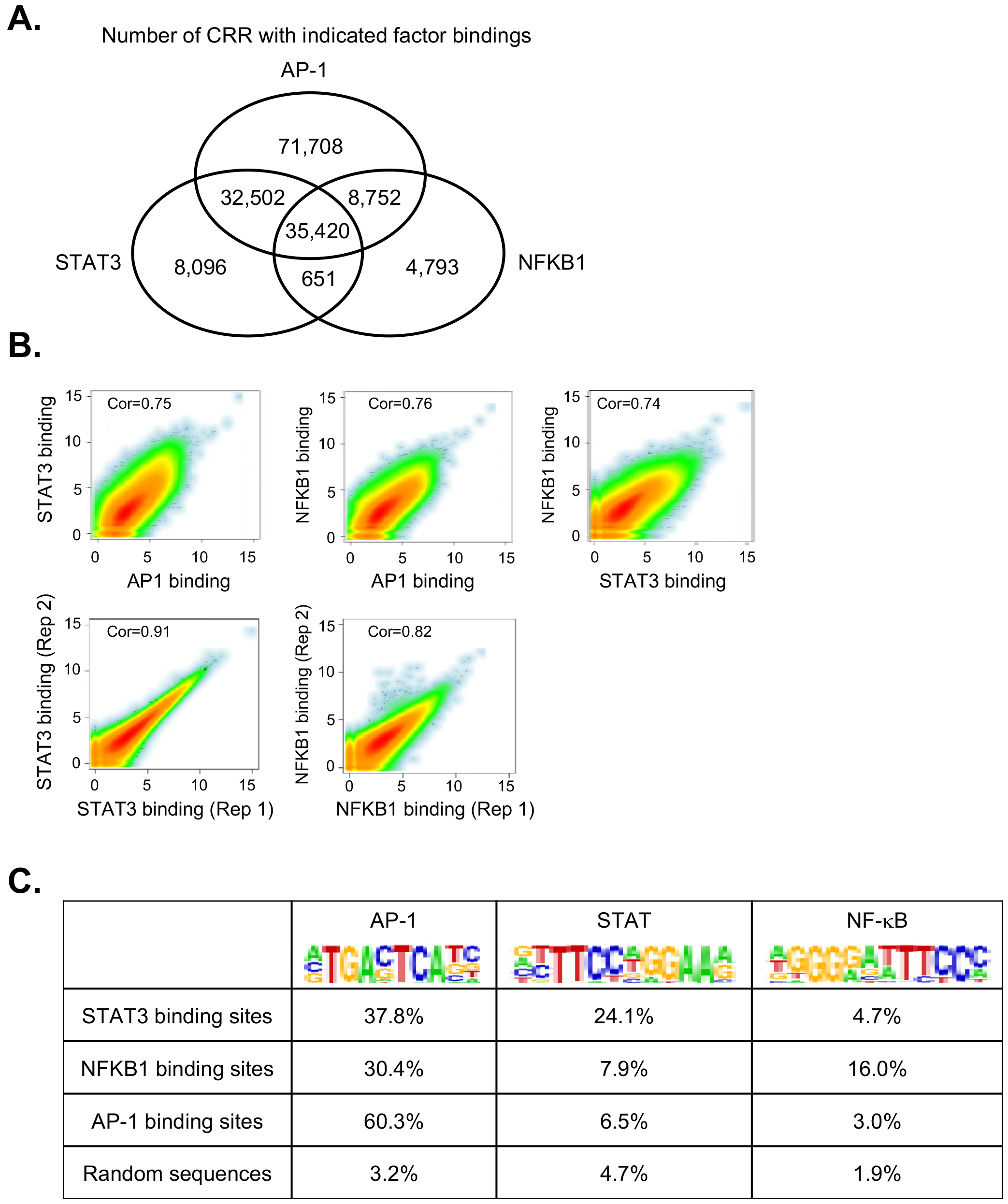
Location analyses and motif analyses of factor binding sites. (A) Number of cis-regulatory regions (CRRs) bound by STAT3, NFKB1 and AP-1. (B) Correlation of AP-1, STAT3, NFKB1 binding levels, and correlation of binding levels between biological replicates. (C) Fractions of factor binding sites with AP-1, STAT, NF-κP motifs. We picked 10,000 random genomic sequences as the control.

**Figure S4.**
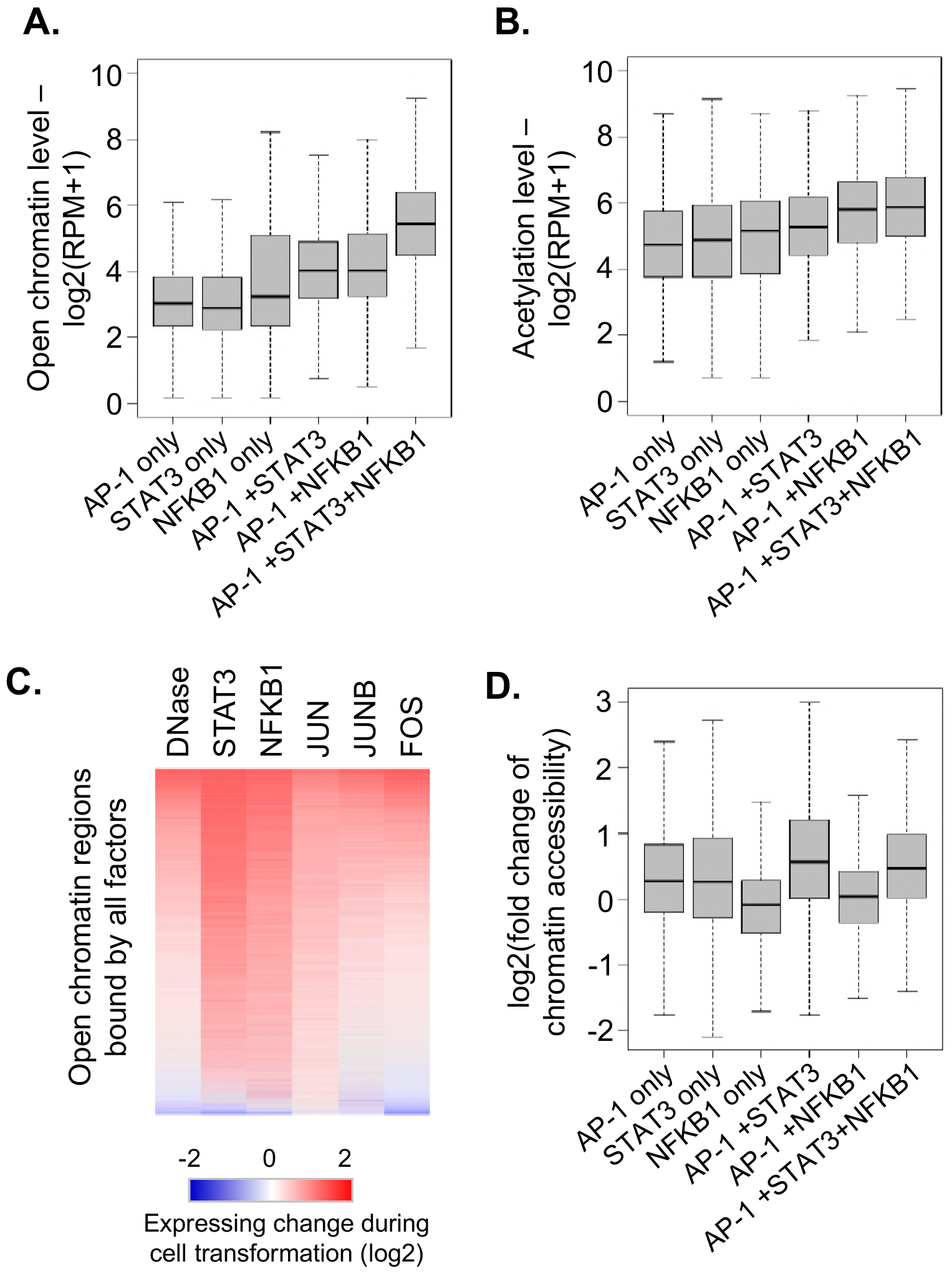
Relationship between factor binding and chromatin accessibility. (A) Chromatin accessibility (DNase-seq) of regions bound by the indicated factors. (B) Histone acetylation (H3K27ac) levels of regions bound by the indicated factors. (C) Heatmap showing the dynamic regulation of chromatin accessibility and factor binding levels for open chromatin regions with binding of the indicated factors. (D) Differential chromatin accessibilities bound by the indicated factors during transformation.

**Figure S5.**
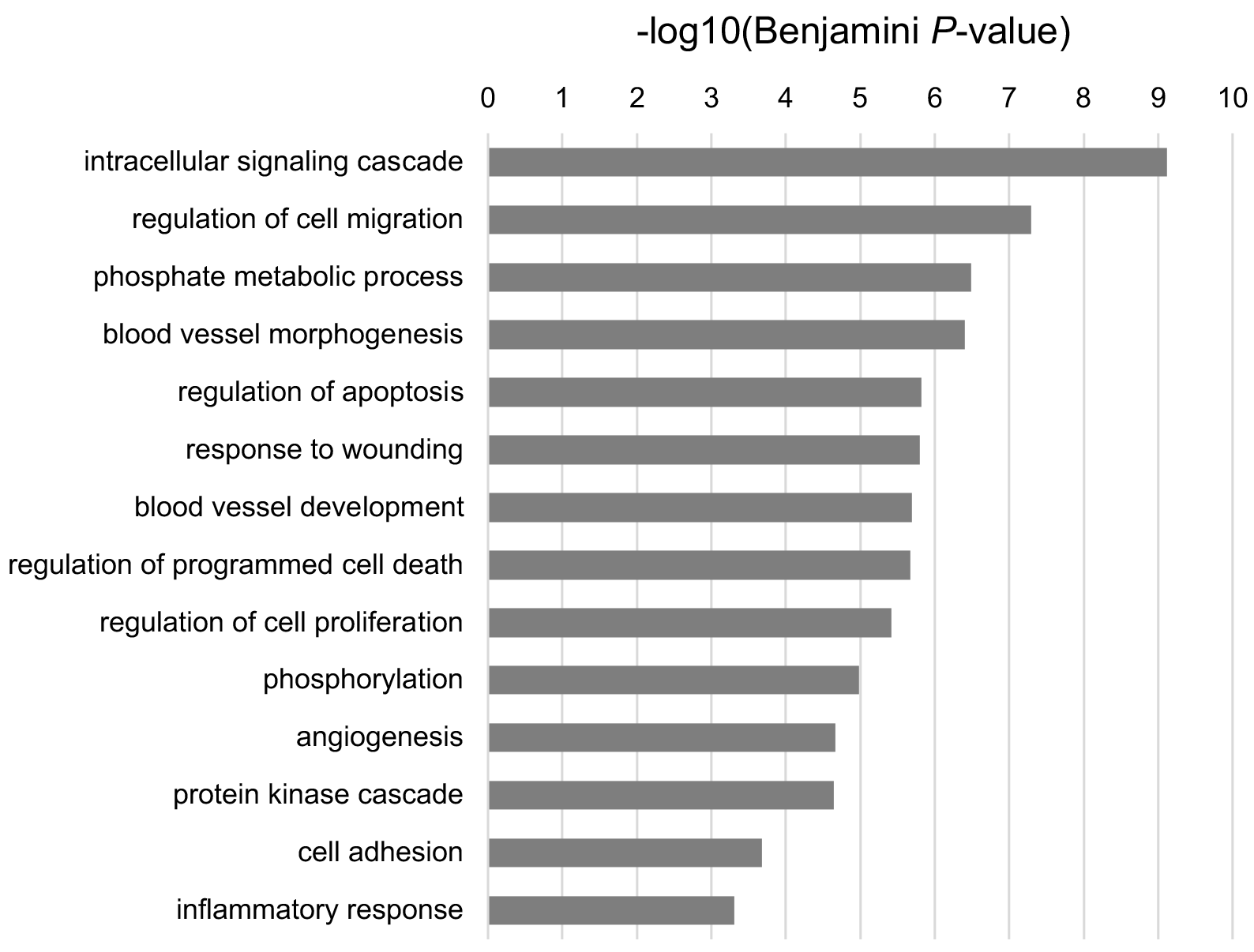
Gene ontology analyses. Classes of genes that are common targets of STAT3, NFKB1, JUN, JUNB and FOS, show >1.5 fold increased binding of at least 4 factors during transformation, and are down-regulated upon at least 4 factor knockouts

**Figure S6.**
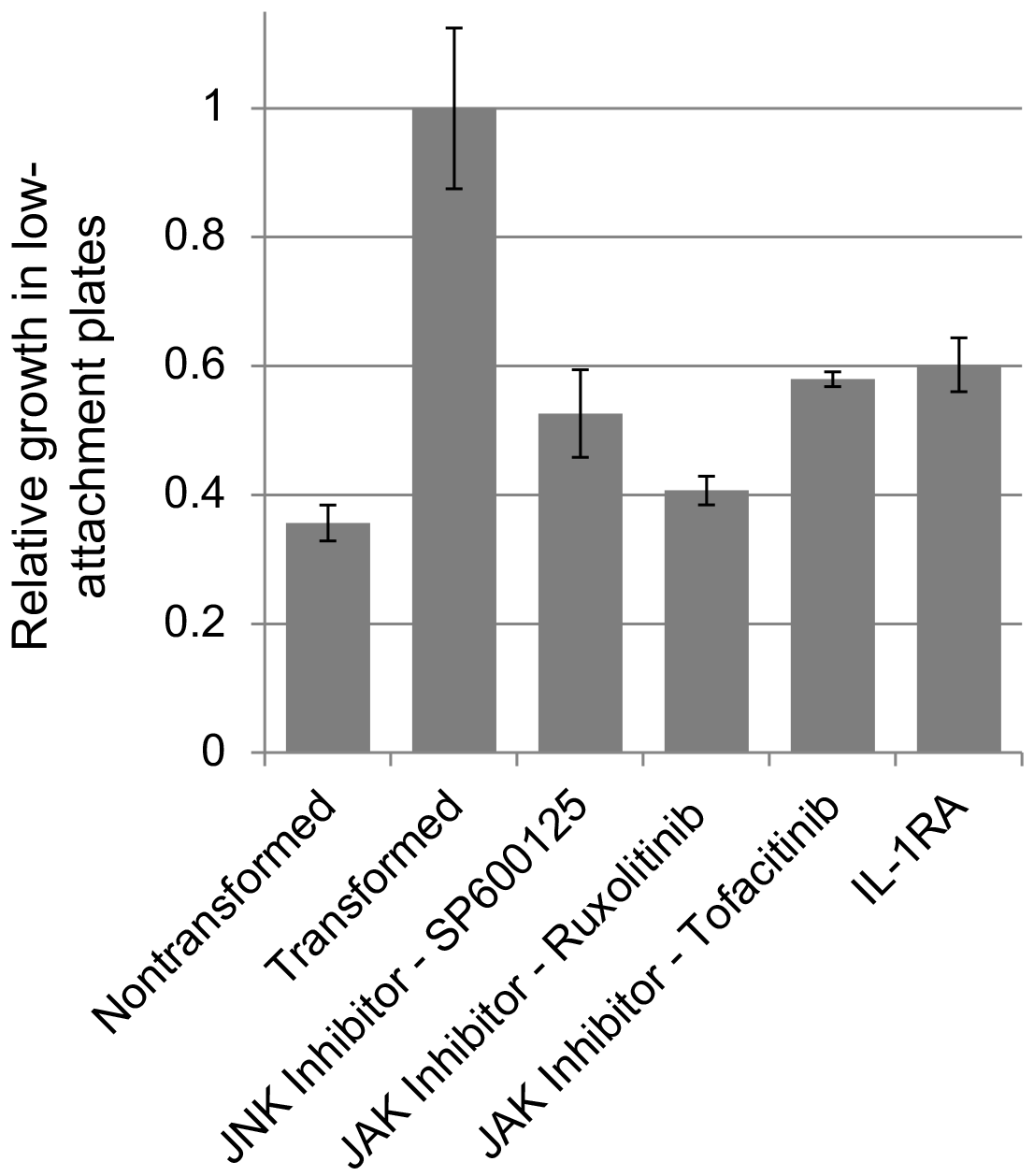
Pharmacological inhibition of JNK, JAK, and IL1 receptor pathways reduces transformation. Transformation efficiencies of cells treated with upon treatment of JNK inhibitor (− 0.3 μM SP600125, 0.3 μM Ruxolitinib, 0.1 μM Tofacitinib, and 30 ng/ml IL1 receptor antagonist.

**Figure S7.**
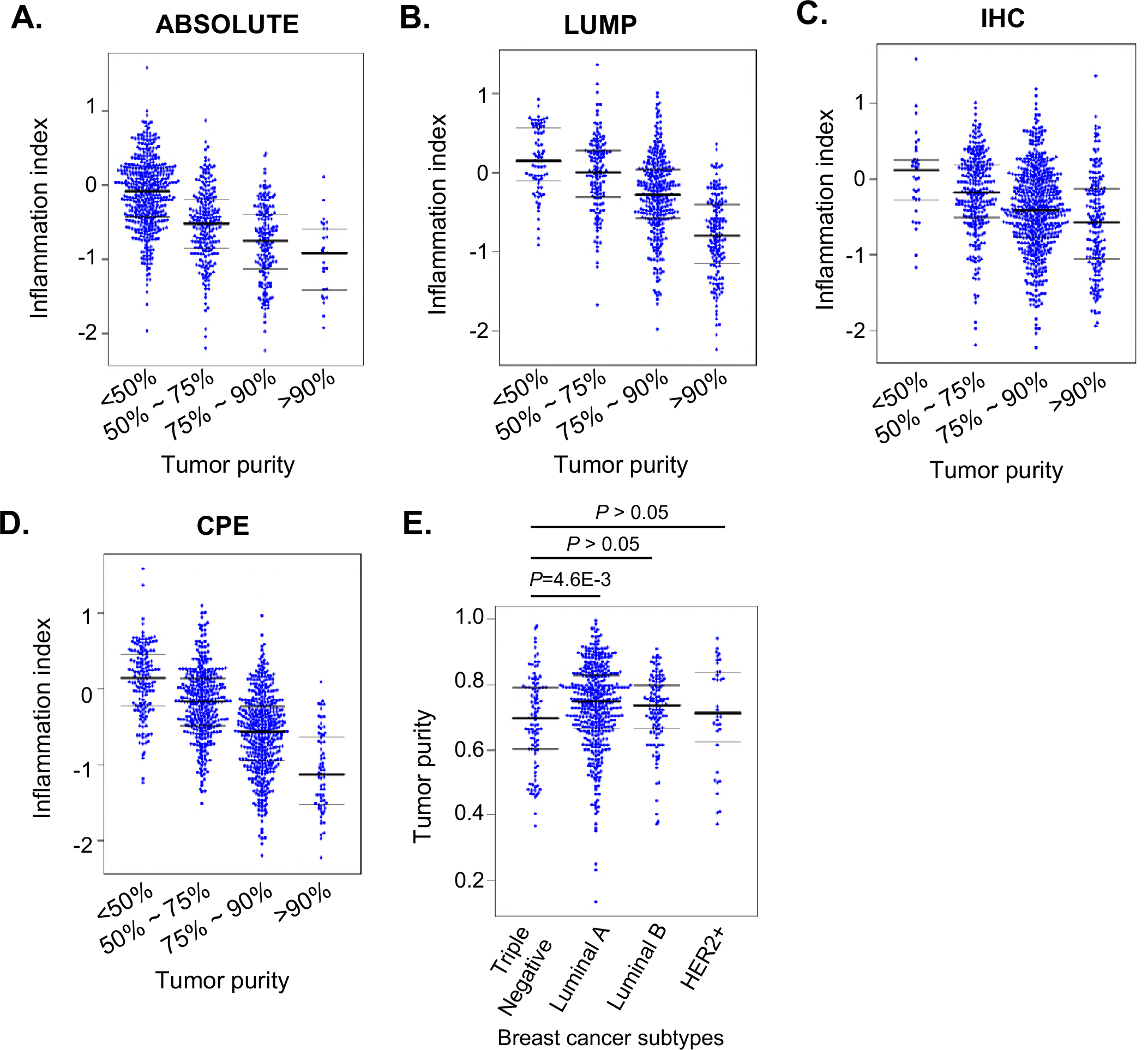
Distribution of the inflammation levels and purity of breast tumors. (A-D) Correlation between tumor purities and the inflammation levels. The tumor purities were estimated using (A) ABSOLUTE, (B) LUMP, (C) IHC, and (D) CPE methods. (E) Purity values of breast cancer subtypes.

**Figure S8.**
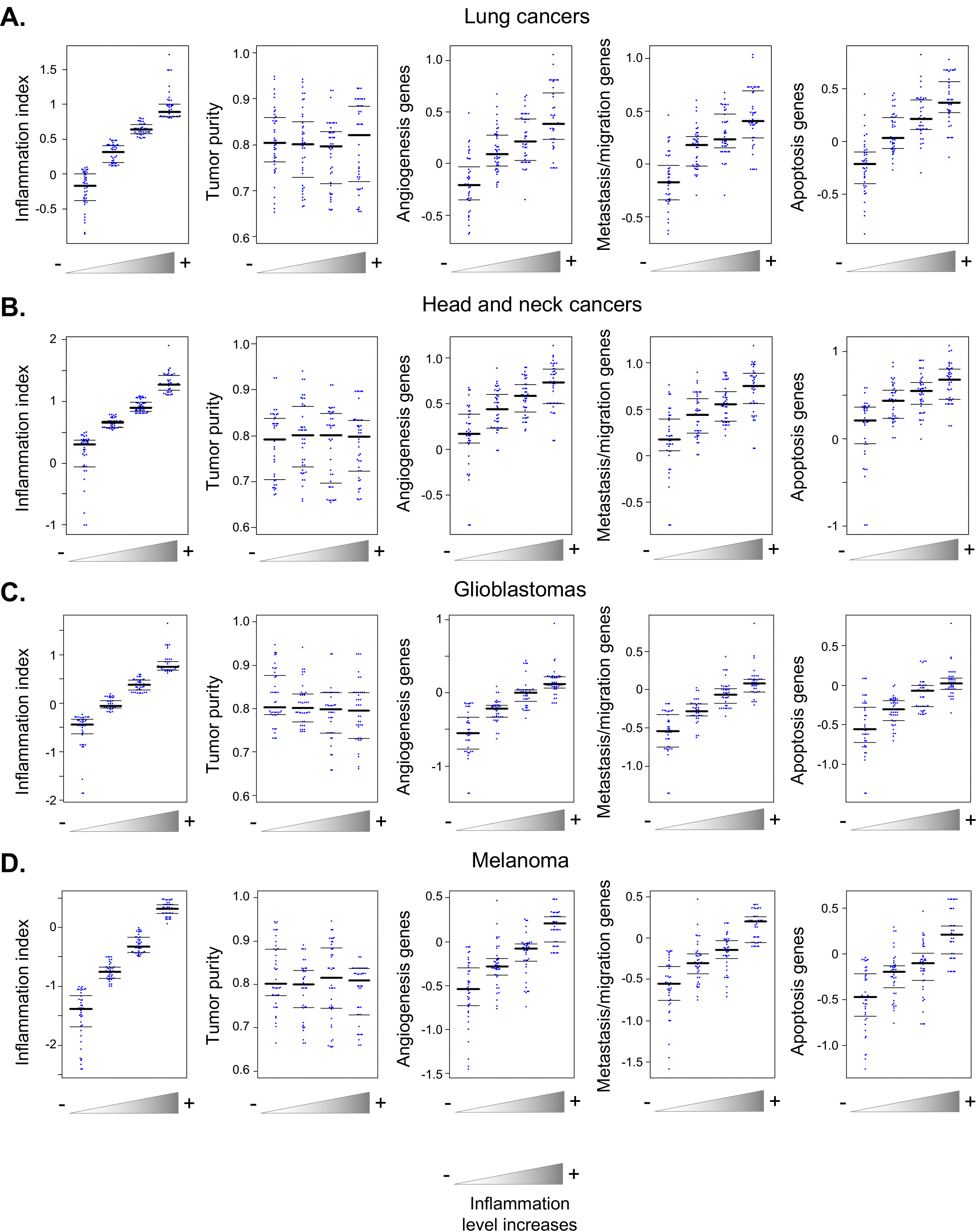
Relationship between tumor purity and inflammation. Randomly picked tumor samples with different inflammation levels and similar purity were examined for expression levels of genes in the indicated oncogenic pathways. Genes related to immune/inflammatory responses were excluded from the analyses.

**Table S1. The direct 1,461 target genes of STAT3/NF-κb and AP1 as defined in Figure 2**

**Table S2. List of sgRNA oligos to knock out the factors.**

